# Juvenile Gouldian Finches (*Erythrura gouldiae*) form sibling sub-groups during social integration

**DOI:** 10.1101/2021.07.16.452682

**Authors:** Gregory M. Kohn, M. Ryan Nugent, Xzavier Dail

## Abstract

The formation of social relationships in complex groups is critical in shaping patterns of social organization and behavioral development. In many birds, young individuals remain dependent on their parents for extended periods but must abruptly transition to navigating interactions in the wider group after independence. While lack of social relationships during this period is detrimental in the development of later social skills, little is known about the social relationship’s juveniles form after independence in many bird species. In this study we describe patterns of social interactions in juvenile Gouldian Finches (*Erythrura gouldiae*) during transitions from family groups into flocks of unfamiliar individuals. A group of 20 juveniles from 4 families were introduced into two flocks. After introductions juveniles showed a gradient of approach rates with most approaches directed towards siblings, followed by juvenile peers, adult females, and lastly adult males. Significant preferences for siblings resulted in the emergence of sibling subgroups within the larger social network. This active self-assortment of siblings suggests that sibling sub-groups are an important bridge linking social connections within the family and the wider group. Such findings suggests that sibling relationships have a critical role in the socialization after independence, as well as structuring the social organization of Gouldian finch flocks.

## INTRODUCTION

Social integration involves establishing relationships in new social contexts. Across the vertebrates, from fish, to birds, to mammals, young individuals remain dependent on their parents for extended periods after birth and switch to interacting with unfamiliar conspecifics during adolescence. In many bird species this switch is abrupt, with juveniles suddenly being cut off from parental support (Collias, 1952; Stamps, Kus, Clark, & Arrowood, 1990). This sudden transition to independence is an important developmental period as it’s associated with high levels of mortality (Lack, 1954) and interactions during this time are necessary for learning to navigate later social challenges (M. West, King, & Kohn, 2011).

Integrating into a new group carries considerable risks and benefits. Being a new group member is associated with having low social status and higher frequencies of aggression from established group members (Marler, 1976; Senar, Camerino, & Metcalfe, 1990). Such risks may be magnified in juveniles who don’t yet possess the necessary skills or experiences to interact with unfamiliar conspecifics. Nonetheless, social integration is a critical component of behavioral development, as juveniles isolated from interactions with others, especially during adolescence, exhibit behavioral and biological deficiencies that impair their ability to thrive in the group (Preiss & Franck, 1974). Therefore, understanding the types of social connections juveniles form as they enter new groups may reveal how they cope with the challenges of social integration.

Many factors influence the formation of juvenile social connections upon entering a group for the first time. Some of these can be passive, caused by constraints in the social or physical environment that force juveniles into close proximity. Passive processes suggest that social connections do not reflect the ability to recognize and prefer specific individuals. For example, in some bird species adults corral their offspring into all-juvenile groups for protection from predation (Munro & Bédard, 1977; Ralf Wanker, Bernate, & Franck, 1996). Such crèches are commonly seen in colonially nesting seabirds and other species, where the risk of predation is high, and juveniles derive substantial protection through the amalgamation of offspring from numerous families. In other cases, juveniles show distinct morphological and physiological differences from adults, which causes them to passively cluster based on shared needs and habitat preferences. For Instance, due to their reduced flying ability fledglings converge on similar spaces as they are unable to synchronize their activities with older individuals. For example, in Piñon Jays (*Gymnorhinus cyanocephalus*), fledglings initially cluster on the ground right after fledging as their lack of flying abilities makes it difficult for them to follow their parents (Balda & Balda, 1978).

Alongside these passive factors, juveniles can form social connections through an active process. In contrast to passive processes active ones involve the recognition and preference for interacting with specific individuals. Upon entering the group, juveniles are exposed to a wide range of potential interaction partners. In groups with a mixed age and sex composition, the choice between interacting with unfamiliar adults, juveniles or siblings may be the first active social decision juveniles make. Numerous studies have shown that birds who inhabit flocks not characterized by strong family ties still possess the ability to recognize siblings and similar-aged peers (Nakagawa & Waas, 2004). Studies on both adults and dependent young (fledglings still being cared for by parents) have shown that birds can differentiate between siblings and unrelated individuals. Adult zebra finch (*Taeniopygia guttata*) males prefer to perch closer to male siblings (Burley, Minor, & Strachan, 1990), and juvenile Greylag Geese (*Anser anser*) can be trained to peck at a specific geometric symbol when in close proximity to particular siblings (Scheiber, Hohnstein, Kotrschal, & Weiß, 2011).

While such studies show that birds are capable of sibling recognition, we know remarkably little about the social connections juveniles make as they integrate into new groups. The evidence we have suggests that sibling relationships provide a bridge between the family group and integration into the wider flock. In Spectacled Parrotlets (*Forpus conspicillatus*), juveniles are herded into crèches by their parents right after fledgling. Here they form strong relationships with siblings that are used as a stable social position to explore interactions with unrelated group memebers (Ralf Wanker et al., 1996). In house sparrows (*Passer domesticus*), individuals follow siblings over non-siblings (Tóth et al., 2009), and in budgerigars (*Melosittacus undulatus*) fledglings attract more interaction from their siblings than expected, and often form an exclusive bond with one specific sibling (Stamps et al., 1990). Cockatiels (*Nymphicus hollandicus*) are more likely to share food with siblings than non-siblings (Liévin-Bazin et al., 2019), and movement and foraging patterns in meadow pipits (*Anthus partensis*), Ospreys (*Pandion haliaetus*) and lilac-crowned amazons (*Amazona finschi*), suggest siblings stay in contact well into independence (Edwards, 1989; Hötker, 1982; Salinas-Melgoza & Renton, 2007).

This study aims to describe the social relationships of newly independent juvenile Gouldian finches during their first transition to a novel flock. Gouldian finches are an endangered estrildid finch that inhabits the subtropical grasslands of northern Australia and exhibit a color polymorphism. Adults show discrete differences in head coloration with some having either red, black, or yellow heads, while juveniles have grey plumage with olive backs. Differences in head coloration correspond with differences in behavior and personality (O’Reilly, Hofmann & Mettke-Hofmann, 2019; Williams, King, & Hoffman 2012; Mettke-Hofmann 2012; Fragueira & Beaulieu, 2019; Pryke, 2007). While little is known about the social structure of Gouldian finches, large fission-fusion flocks composed of hundreds of individuals in mixed age and sex composition are commonly observed in the wild. The fluidity and diversity of estrildid flocks has led some to speculate that they lack significant patterns of within-group structure (Bolton et al., 2016; Dostine et al., 2001).

The social relationships individuals form will directly shape the organization and experience of living in a group (Aureli & Schino, 2019). By describing how juvenile Gouldian finches integrate into novel flocks, this study aims to uncover significant aspects of this species developmental ecology. By identifying which social relationships are preferred and avoided during integration, we can uncover if juveniles are preferentially exposed to influences from specific group members during this critical period. In the wild, small aggregations of juvenile Gouldian finches are observed around waterholes, with adults descending to the waterhole first, followed by groups of juveniles (O’Reilly, Hofmann, & Mettke-Hofmann, 2019). However, it’s unknown if juveniles maintain connections with peers and siblings outside the waterhole, or if associations between juveniles at these locations reflect sibling or peer relationships. Due to the presence of sibling recognition abilities in related species (Nakagawa & Waas, 2004) we hypothesize that newly independent juvenile Gouldian finches will maintain strong relationships with siblings upon entering the flock for the first time.

## METHODS

### Subjects

All individuals in this study were Gouldian finches. All finches were acquired from professional breeders in Arizona and Florida, U.S.A. Each bird was provided with a uniquely colored leg band for individual recognition. Males, females, and juveniles were distinguished from each other based on plumage and behavioral differences, with adult females being much more subdued in their chest coloration in contrast to males.

All juveniles were hatched and raised in separate cages prior to being introduced to experimental flocks. At around 40 days after fledging, juveniles become independent of their parents and can feed and drink on their own without any sign of parental provisioning. Juveniles were distinguished from adults based on their grey heads, olive backs, and lack of typical adult plumage. In this study, all juveniles were under one year of age since fledging the nest and adults were older than a year and exhibited adult plumage. Head color was easily observed in adults and assessed in juveniles at the time of their first molt after the end of the data collection period. We used a total of 11 adults and 20 juveniles for this study.

All birds were provided with a daily diet consisting of finch seed mix (Kaytee), with soaked Perle Morbide pellets, eggshell, boiled eggs, and dandelion greens provided twice a week. Fresh water was continually available.

### Aviaries

We used a series of two aviaries in this study. Each aviary consisted of identical dimensions (23 x 72 x 41 in) and similar environmental conditions. The aviaries were located indoors next to a large window allowing for natural sunlight and ventilation. Each aviary contained both natural eucalyptus and plastic privacy perches, small shrubs, a feeding station, and a bowl for water baths.

### Data Collection

We used focal sampling during data collection. During observation periods, all individuals in the flock were observed for a 5-minute focal block. We recorded all approaches that occurred within a 2-inch radius around the focal individual being observed. All observations were done using voice recognition technology allowing the observer to maintain continual visual contact with the focal bird throughout the sampling block. An approach was scored when another individual approached another within a radius of 2 inches. Before data collection, video of marked perches was made to establish that 2 inches was the typical social distance observed between two resting adult finches. All focal samples were collected by GMK.

#### Procedure

##### First introduction

On 10/5/2020, four novel juvenile siblings were introduced into an aviary consisting of 6 adult female and 5 adult males. From 10/06/2020 to 12/22/2020, 503 focal samples were taken. All individuals observed for 34 blocks for a total of 170 minutes except one individual who was removed from the flock temporarily for health reasons and was observed for 26 blocks.

##### Second Introduction

On 2/10/2021, the aviary from the first introduction was split up into two separate flocks: three adult males, two females, and two juveniles were moved into Aviary 1, while four females, two males, and one juvenile were moved to Aviary 2. In each flock, three new juvenile siblings were randomly introduced. From 2/11/2021 to 4/20/2021, a total of 84 blocks were recorded in each aviary. Everyone was observed for a total of 8 blocks measuring 40 minutes each.

##### Third Introduction

On 05/25/2021, a group of 5 juveniles were transferred from their family group and introduced into Aviary 1. On 06/11/2021, another group of 5 siblings were transferred to Aviary 2. From 05/26/2021 to 06/27/2021, all individuals in the flock were focal sampled, collecting a total of 240 blocks in each aviary. Each individual was observed for 16 blocks for a total of 80 minutes each.

## ANALYSIS

We used generalized linear mixed models to investigate the social preferences of juveniles when integrating into the flock. Three models were constructed for this study: the dyad-model, age-model, and sibling-model. The dyad-model was a multi-membership model performed using the MCMCglmm package and was used to predict the dyad weights (how often two individuals in the flock interacted) for all dyads involving juveniles. The MCMCglmm package is a Bayesian mixed model package that allows for multi-membership and random effects (Hadfield, 2010), and has been used to investigate the factors shaping social networks and relationships in animals (Hart, Weiss, Brent, & Franks, 2021; Raulo et al., 2021). Trace plots were examined to assess if the dyad-model was well-mixed, and Heidelberger and Welch’s diagnostics were used to test if the Markov chain was from a stationary distribution. The dyad weights represented the number of times each dyad in the flock approached each other. For each dyad we classified if it was between members of the same sex, age, family, or head color as a fixed effect. The random effects included the effect of round, aviary, and a multi-membership term that contained the initiator and receiver of each dyad.

Post-hoc tests included a permutation test where raw approach data was permuted 100,000 times to create a null distribution of approaches for each aviary. The observed approach rates were then compared to the permuted approach rates to see if they occurred above or below null expectations with empirical p-values calculated at an alpha of 0.025 for each tail of the distribution.

The second model, the age-model investigated if juveniles showed preferences towards adults or juveniles in the aviaries. The third model, the “sibling-model”, investigated if juveniles showed preferences towards siblings and unrelated individuals. Both models were generalized linear mixed models conducted using the lmerTest package (Kuznetsova, Brockhoff, & Christensen, 2017). We first calculated the rate of approaches towards juvenile, adult male, adult female, sibling and non-sibling conspecifics. This was done by dividing the sum of the approaches toward members of a specific category, such as juveniles, by the number of individuals in that category in the flock. Both models included the approach rate as the dependent factor. The age-model included approach rates towards adult males, adult females, and juveniles, while the sibling-model included the approach rates for siblings and unrelated flock members. Each model included the direction of approach’s (age-model: adult male, adult females, or juveniles; sibling-model: siblings or non-siblings), the individual’s sex, and head color as fixed effects with random effects of individual identity, round, and aviary. Post hoc tests used Wilcoxon tests to run pairwise comparisons between interaction preferences while controlling for multiple comparisons using the False Discovery Index.

## RESULTS

During the first introduction a group of four juveniles was added to the aviary and a total number of 8,840 approaches were recorded. In the second introduction the flock was split up into two separate aviaries and three juveniles added to each aviary. Here a total of 2,184 approaches were recorded in Aviary 1 and 2,345 were recorded in Aviary 2. During the third introduction where 5 juveniles were introduced to each aviary a total of 4,425 approaches were recorded in Aviary 1 while 3,990 approaches were recorded in Aviary 2.

The dyad-model showed a significant influence of family, age, and head color on the frequency of dyads. The strongest dyads involving juveniles were more likely to occur between members of the same family and between individuals of the same age. Our model also uncovered a significant effect of head color with the coefficient indicating stronger dyads for individuals of differing head colors (Table 1). Post-hoc permutation analysis showed that, across all the flocks, the juvenile-to-juvenile dyads occurred at a higher rate than expected under null conditions, while in many introductions the juvenile-to-male dyads occurred at lower-than-expected rates (Table 2).

**Table 1:** Results from the Dyad Model Results from the Monte-Carlo Markov Chain Generalized Linear mixed model. The effect of same sex, age, and family was included in the model as fixed effect. The

| Fixed Effects | Posterior means | Lower 95% CI | Upper 95% CI | Bayesian *P* value |
| --- | --- | --- | --- | --- |
| **Sex** | 2.76 | -0.8377 | 6.3081 | 0.64 |
| **Family** | 30.39 | 25.5280 | 34.9899 | <0.001 |
| **Age** | 10.88 | 5.8812 | 16.1538 | <0.001 |
| **Head Color** | -7.17 | -11.4511 | -3.1861 | <0.001 |

**Table 2:**
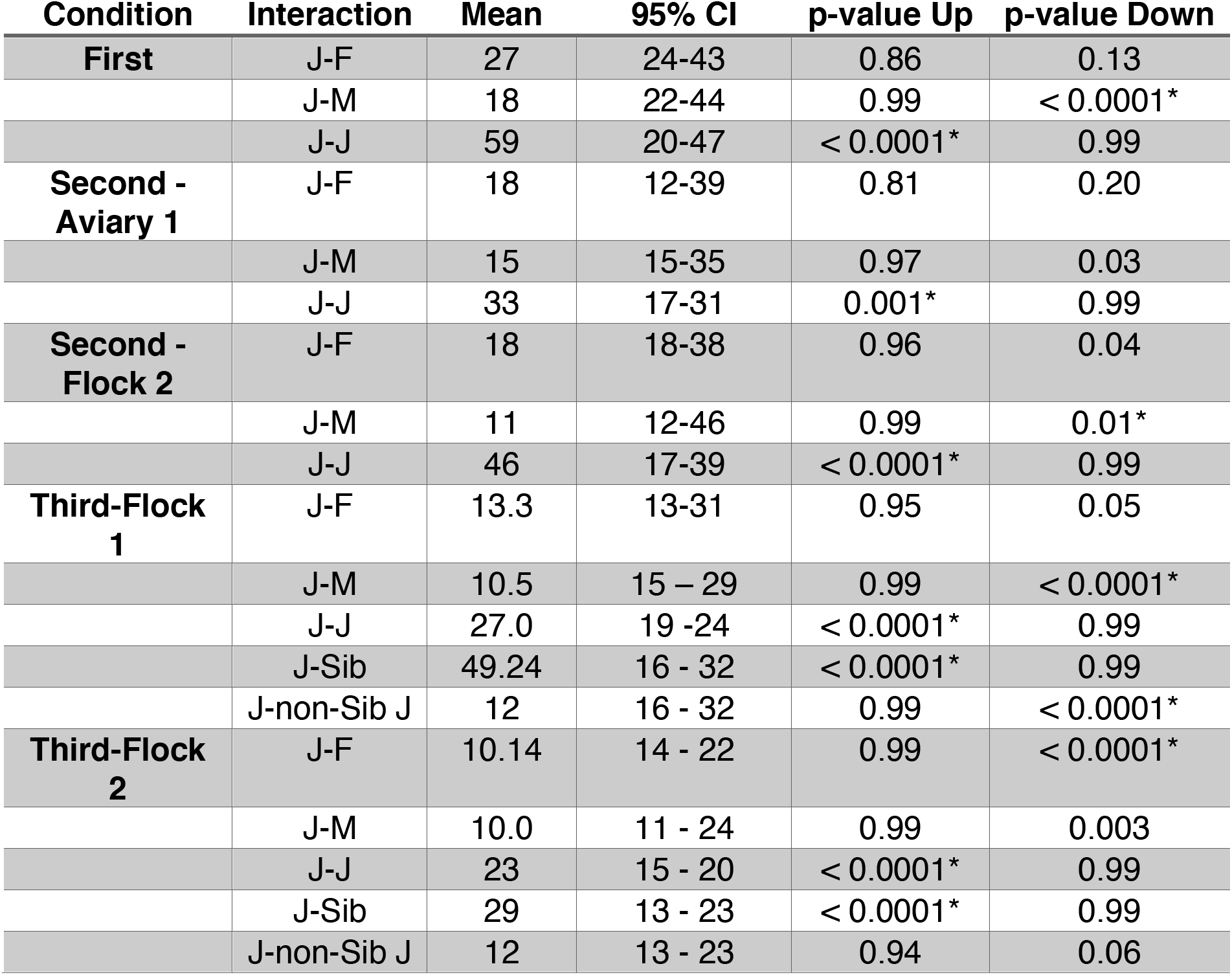
Permutation Tests for Approaches by Age and Sex

Both the age and sibling model showed a significant main effect of approach direction. The age-model showed a significant main effect of approach direction toward juveniles or adults (Table 3) and the sibling-model showed a significant main effect of approach direction towards siblings and non-siblings (Table 3). Post hoc tests indicated juveniles had higher approach rates towards other juveniles than towards adult males and females (N = 20, Mean juvenile approach rate = 40.65, Mean female approach rate = 18.01, Mean male approach rate = 11.83, Wilcoxon Rank Sum test, juvenile-adult female, V = 0, *p* < 0.0001, Wilcoxon Rank Sum test, juvenile-adult male, V = 0, *p* < 0.0001, Fig 1). When interacting with adults, juveniles showed higher rates of approaches towards females in contrast to adult males (N = 20, Wilcoxon Rank Sum test, adult male-adult female, V = 26, *p* = 0.005). Approach rates towards siblings where higher than approach rates to unrelated conspecifics (N = 20, Mean sibling approach rate = 23.6, Mean non-sibling approach rate = 14.15, Wilcoxon Rank Sum sibling-non-sibling, V = 163, *p* = 0.03172, Fig 1).

**Table 3:** Results from the age, sibling, and sibling-juvenile model

| Model | Fixed-Effect | Coefficient | Std. Error | t value | P value |
| --- | --- | --- | --- | --- | --- |
| **Age** | Direction | 22.556 | 2.559 | 8.813 | <0.001* |
|  | Sex | -5.817 | 4.862 | -1.196 | 0.2455 |
|  | Head Color | 2.641 | 2.314 | 1.141 | 0.2673 |
| **Sibling** | Direction | 9.4500 | 2.8992 | 3.260 | 0.0023* |
|  | Sex | 6.1024 | 3.7654 | 1.621 | 0.12 |
|  | Head Color | 2.3855 | 5.2048 | 0.458 | 0.68 |
| **Sibling-Juvenile** | Direction | 6.3000 | 1.9893 | 1.146 | 0.005* |
|  | Sex | 2.7647 | 2.4123 | 1.146 | 0.27 |
|  | Head Color | 7.0147 | 4.0366 | 1.543 | 0.35 |

**Figure 1:**
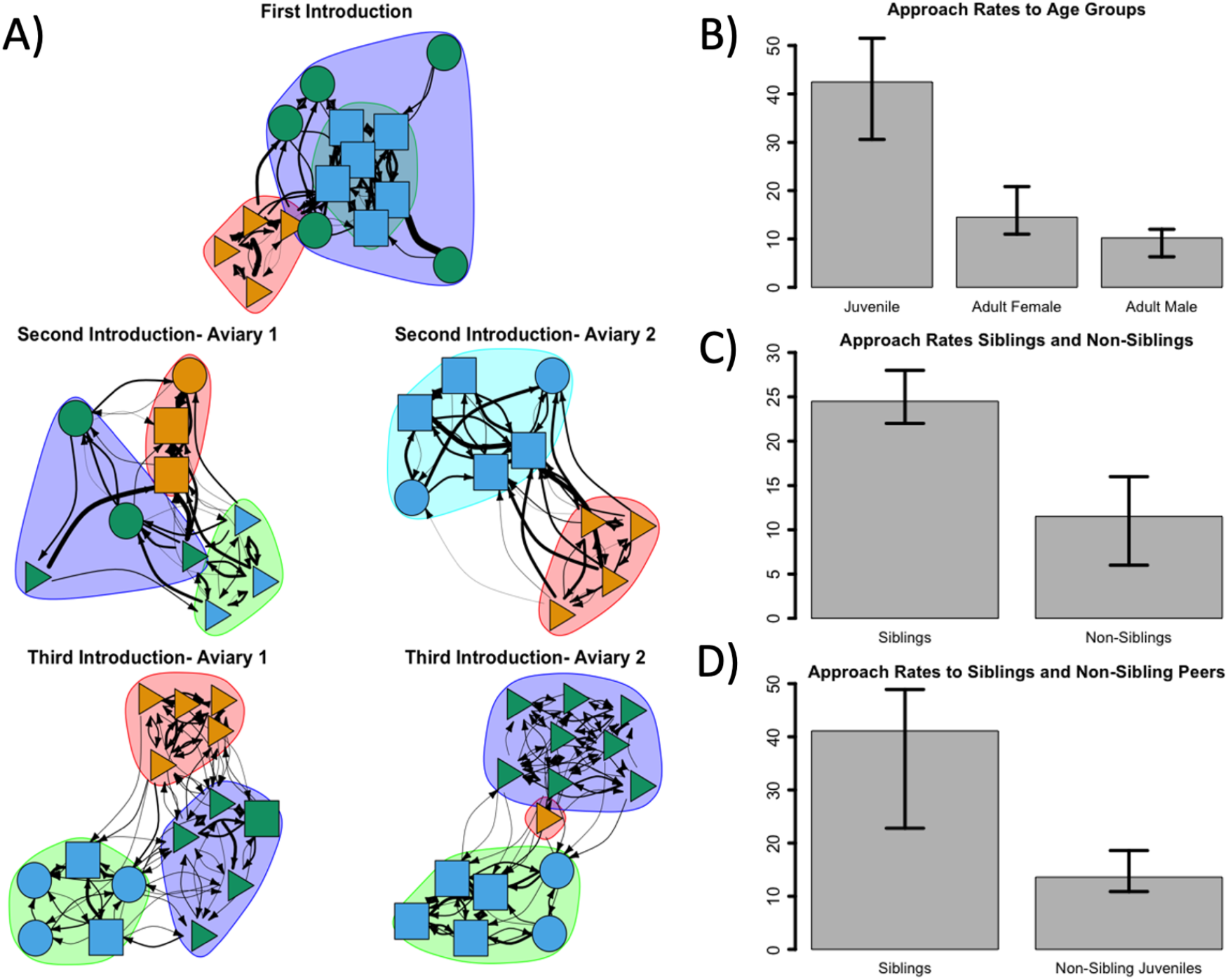
Social Networks and Social Preferences A) Social Networks formed during the different introductions. Shapes represent individuals, with triangles as juveniles, circles as adult males and squares as adult females. Arrows represent approaches between individuals. The spinglass communities are shown in the grouping of different shapes. B) Bar graph of the median approaches directed towards juveniles, adult females and adult males. The error bars represent 95% confidence intervals in all graphs. C) Bar graphs represent median approaches directed towards siblings and non-siblings. D) Bar graphs represent median approaches siblings and non-sibling juveniles during the third introduction.

To see if juveniles preferred related siblings over unrelated juveniles, we did an additional analysis restricted to our third introduction. During the third introduction, the flock size was large enough to calculate approach rates towards both siblings and unrelated juveniles for the 10 juveniles introduced during this period. An additional GLMM was performed using the lmerTest package that looked at the fixed effect of direction (sibling versus non-sibling juveniles), sex, and head color with random effects of aviary and individual. Our model showed a significant influence of direction (Table 3) with the juveniles preferring to interact with siblings over unrelated peers (N = 10, Mean Sibling = 39.02, Mean unrelated juvenile = 16.7,, Wilcoxon Rank Sum test, siblings-unrelated juveniles, V = 55, *p* = 0.002, Fig 1 C). Permutating approaches during the third period indicated that while sibling directed approaches occurred above null expectations, approaches to unrelated juveniles occurred below null expectations (Table 2).

## DISCUSSION

This study described the social preferences and relationships of newly independent Gouldian finches during their transition from the family group to the larger flock. The period between fledging and adulthood is one of the most critical stages for survival and development in birds. Our results show that juvenile Gouldian finches preferentially choose to interact with siblings after becoming independent of their parents. This is among the few examples of sibling-assortment of independent juveniles in passerine flocks (Hötker, 1982; Kohn, King, Scherschel, & West, 2011; Nicolai, 1956; Templeton, Reed, Campbell, & Beecher, 2012; Tóth et al., 2009). Upon entering the group for the first time, juveniles showed a gradient of approach rates with most approaches directed towards siblings, followed by juvenile peers, adult females, and lastly towards adult males (Fig 2). Juveniles showed significantly higher approach rates towards siblings over both unrelated juveniles and adults. This resulted in the emergence of sub-groups that remained consistent across changes in the age-sex composition of the flock.

**Figure 2:**
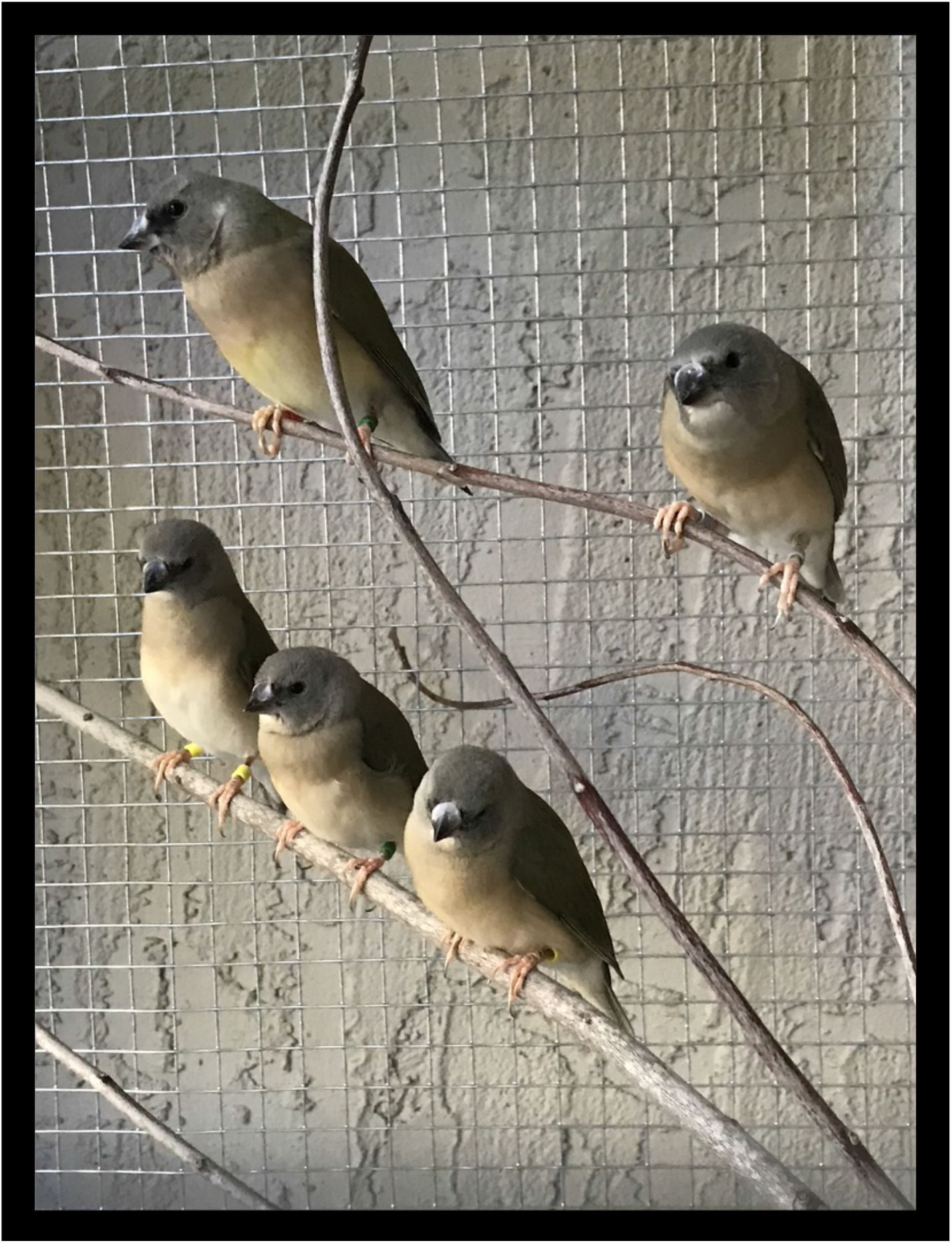
Picture of Juvenile Subgroup Picture of juvenile subgroup. Associations between juveniles often resulted in visually clustered associations within the larger flock.

A diverse range of factors can cause the formation of sibling and peer groups in juvenile birds. In zebra finches, fledglings initially follow their parents around the colony but quickly cluster around areas with same aged-peers (Goodwin, 1982; Ogino, Maldonado-Chaparro, & Farine, 2021; Zann, 1996). In these contexts, it is unclear if the close association between juveniles is caused by preferences to interact with other juveniles, collective herding of juveniles by parents, or a juvenile’s inability to synchronize their behavior with parents during foraging trips. Therefore, understanding the role that the environment, experiences, morphology, and active social preferences play in the formation of juvenile subgroups is essential to understanding their function in the evolution and development of avian behavior.

Our results suggest that the formation of juvenile subgroups in Gouldian flocks is an active process, reflecting the ability of juveniles to recognize their siblings and mutually prefer interacting with them. Studies in birds have shown that individuals can recognize and preferentially attend to siblings (Nakagawa & Waas, 2004). In a cross-fostering study with juvenile Bengalese finches (*Lonchura striata domestica*), preening bouts between genetic siblings were longer and more frequent that preening bouts between foster siblings (Ju & Lee, 2016). In house sparrows (*Passer domesticus*), juveniles preferred to follow siblings (Tóth et al., 2009) over unrelated conspecifics. Our analysis showed that sibling-to-sibling approaches were significantly higher than approaches to all others, and that juvenile subgroups were more likely to contain siblings than unrelated juveniles (Fig 2). In captive contexts, competition over patchy resources is minimized and observers did not find that either juvenile or adults consistently occupied different areas of the enclosure (Kohn, personal observations). Outside of plumage differences, juvenile Gouldian finches do not show substantial morphological or physiological differences from adults (Gelis, 2003). This suggests that the higher approach rates towards siblings observed in juvenile Gouldian finches reflects decisions made in accordance with underlying social preferences.

In the wild, the period from fledging to independence is characterized by high mortality and unique developmental challenges (Cox, Thompson III, Cox, & Faaborg, 2014). Specifically, one of the largest challenges juveniles face is integrating into novel groups for the first time with little experiences outside the family (Collias, 1952). Our study highlights that siblings play an important role in social integration as they are the strongest relationships juveniles have after becoming independent of their parents. The existence of sibling sub-groups assures some social connections will remain stable as juveniles move from the family to the larger flock. This provides a predictable social context that can buffer against aggressive interactions with novel individuals. Interactions with novel conspecifics carries increased risk of aggression since adults are often more aggressive toward unrelated juveniles in birds (Senar et al., 1990). Juveniles may respond to adult aggression by quickly becoming selective in their interactions and preferentially associating with familiar siblings. Sub-groups of familiar siblings will soften the impact of stressful interactions with novel individuals while assuring individuals get the predictable social experiences needed for development.

Interactions with siblings can have significant benefits, such as access to beneficial social information or experiences necessary for the development of competent behavioral abilities (Alberts, 2008). Among mammals and birds, early interactions are critical for the development of species-typical abilities (King & West, 2002; M. J. West & King, 1987). When isolated from peers, juveniles often develop distinct behavioral repertoires or long-lasting behavioral deficiencies when compared to typically reared individuals (Suomi & Harlow, 1972;). For instance, Rosy-faced lovebird (*Agapornis roseicollis*) chicks raised in isolation were more aggressive, showed increased avoidance behavior, were unable to form a pair bond, or become integrated into the flock (Preiss & Franck, 1974). In mallard ducklings (*Anas platyrhynchos*) chicks require direct interactions with siblings in semi-naturalistic environments to properly imprint on maternal cues (R. Lickliter & Gottlieb, 1985). Ducklings will even switch their imprinting preferences to be in line with their siblings if they show a preference for different objects (R. Lickliter & Gottlieb, 1986). In cross fostering studies on zebra finches, it was shown that the ability to imprint on foster Bengalese finch parents is shaped by the presence of siblings. Zebra finches raised by Bengalese finch parents, but with 4 to 5 zebra finch siblings showed species-typical sexual preferences whereas those with fewer siblings developed sexual preferences for Bengalese finches (Kruijt, Ten Cate, & Meeuwissen, 1983). While such results show that siblings are necessary resources in development of species-typical abilities, less is known about the ontogenetic role that siblings play after independence.

Upon independence, behaviors acquired through sibling and peer interactions can facilitate integration into the group (Pellis & Pellis, 2007). Interactions with siblings allows individuals to explore their behavioral potentials and refine their behavioral abilities without the risks associated with adult interactions. In Spectacled parrotlets (*Forpus conspicillatus*), interactions within siblings serve as a springboard to the formation of their first pair bonds (Garnetzke-Stollmann & Franck, 1991; Ralf Wanker et al., 1996), and singletons who hatch without siblings compensate by prolonging their relationship with parents and forming wider social networks at independence (R. Wanker, 1999). In Brown-headed cowbirds (*Molothrus ater*), interactions in juvenile subgroups predict the development of competent courtship behavior during an individual’s first breeding season (Kohn, King, Dohme, Meredith, & West, 2013). Rhesus macaque (*Macaca mulatta*) infants raised in isolation were able to acquire species-normative social behavior after an extensive period of social interaction with peers (Suomi & Harlow, 1972). Some studies have even shown that juveniles cluster to form “gangs” around contested resources to better compete with more experienced adults (Dall & Wright, 2009; Franks et al. 2020). These studies suggest that juvenile and sibling subgroups may facilitate the horizontal co-construction of social skills necessary to navigate interactions during later periods (Kohn, 2019). Current studies are looking at the details of juvenile-to-juvenile interactions in Gouldian finches to explore how they might guide the development of competence social and reproductive skills in adulthood.

When interacting with adults, juvenile Gouldian finches showed a significant preference to interact with females over males. In many species, females are less aggressive towards juveniles in contrast to males (Ward & Webster, 2016). When exploring connections outside sibling subgroups, juveniles experience less displacement and aggression from females in contrast to males (Kohn, in prep). Across the different flocks, juvenile approaches towards males were more likely to occur below null expectations, suggesting that females may be the first unrelated adults juveniles have consistent contact with. While little is currently known about the shifting patterns of social relationships across development in birds this study suggests that adult females may be an important factor in social integration.

The stability of a group’s social organization reflects the ability of juveniles to reconstruct patterns of relationships across generations (Deputte, 2000; Kohn, 2019). Consequently, the social relationships made by juvenile finches as they transition into the flock for the first time will influence how they experience their social environment and the social organization of the group itself. By constructing sibling sub-groups, juvenile Gouldian finches will disproportionately have access to information from siblings. This study and others suggest that sibling relationships are likely an important component in the social development of passerines. As passerines are heavily dependent on socially transmitted information, the emergence of a stable network of siblings may create the predictable social contexts necessary for the development of competent social skills.

## ACKNOWLEDGMENTS

This study was supported with a grant from the Van Vleck Award. All research was done in accordance with ASAB/ABS Guidelines for the Use of Animals in Research and was approved by the University of North Florida’s Institutional Animal Care and Use Committee protocol number 20-006.

